# Rice biological nitrification inhibition efficiency depends on plant genotype exudation rate

**DOI:** 10.1101/2023.05.31.543046

**Authors:** Jasmeet Kaur-Bhambra, Joy Ebenezer Rajakulendran, Dylan Bodington, Marcel Jaspars, Cécile Gubry-Rangin

## Abstract

Nitrification largely contributes to global nitrogen (N) fertiliser loss and nitrous oxide emissions in agricultural soils, including rice cultivation, Asia’s largest fertiliser consumer. One promising mitigation strategy to achieve greener agriculture involves biological nitrification inhibition (BNI) by plant-derived compounds. Future implementation of this nature-based approach in agricultural settings requires a better understanding of the impact of plant physiological traits on BNI efficiency and nitrification dynamics. We targeted those objectives in five rice genotypes grown in greenhouse conditions. The BNI efficiency was variable among the five plant genotypes, with a stronger inhibition of the ammonia-oxidiser in the rhizosphere than in the bulk soil. We identified that the root mass, root exudation rate and chemical composition are factors explaining the distinct BNI efficiencies in the rice genotypes, with plants having a high BNI efficiency having a small root mass and a high root exudation rate. Using the BNI efficiency assay of root exudates on multiple AO cultures, we demonstrated that AO bioassay could accurately represent the BNI variability in the soil. Finally, we identified a novel BNI compound, *N*-butyldodecane-1-amine (NBDA), in two high-BNI genotypes. NBDA specifically inhibited ammonia oxidisers by inhibiting enzymes involved in the ammonia oxidation pathway. These findings demonstrate that BNI research integrating plant physiology, microbial ecology, and chemistry has a strong potential for providing more sustainable agriculture.

## Introduction

Agricultural soils, which require over 150 mT Nitrogen (N) fertiliser per year (Ladha et al., 2016; FAO., 2019), are the source of approximately 65% of nitrous oxide emissions (IPCC 2007), which is strongly impacted by nitrification by consuming up to 50% of the applied fertilisers (Cassman et al., 2002). Soil nitrification can be inhibited by some plant root exudates, a mechanism termed Biological Nitrification Inhibition (BNI) (Subbarao et al., 2009). As nitrifiers compete for N with plants, BNI has strong potential to increase the nitrogen use efficiency (NUE), especially in highly fertilised crops, such as wheat (Subbarao et al., 2007), sorghum (Zakir et al., 2008), maize (Otaka et al., 2022) and rice (Tanaka et al., 2010).

Rice (Oryza sativa L.) is a worldwide staple food for half of the global population, and its production is crucial for world food security (bin Rahman & Zhang, 2022). To meet its ever-growing demand, the agricultural rice sector accounts for approximately 16% of the total N fertiliser usage (Ladha et al., 2016), resulting in 6.8 Tg of N_2_O emissions per year (IPCC 2007). Difference in NUE and nitrification activity was demonstrated in some rice cultivars (Chen et al., 2022; Li et al., 2007). This variability was explained by a range of BNI efficiency in 36 rice genotypes through testing growth inhibition of *N. europaea* AOB by the rice root exudates (Tanaka et al., 2010). BNI efficiency of some rice cultivars was not only demonstrated in hydroponic conditions but also a range of soils (Chen et al., 2022; Lu et al., 2022; Illarze et al., 2021; Sun et al., 2016). Two structurally different BNI compounds, a fatty alcohol, 1,9-decanediol (Sun et al., 2016) and a phenolic compound, syringic acid (Lu et al., 2022), have been isolated from rice root exudates, even showing some synergetic effects (Lu et al., 2022). The BNI compound 1,9-decanediol was assessed as an efficient BNI compound on a range of AOA and AOB cultures, with higher sensitivity for AOA than AOB (Kaur-Bhambra et al., 2022). The composition of root exudates is complex (Dietz et al. 2020), with 1, 9-decanediol representing only 36% of rice BNI exudates (Sun et al., 2016), suggesting the presence of other BNI compounds and potential synergetic effects (Lu et al., 2022).

These advances in understanding rice BNI are supported by multidisciplinary research efforts in other crops, including developments on plant genetics (Subbarao et al., 2021), biochemistry (Egenolf et al., 2020), soil science (Lu et al., 2019), microbiology (Nardi et al., 2020), agronomy (Subbarao & Searchinger, 2021), and panomics (Ghatak et al., 2023). This later field holds some strong promises to address many research gaps to understand the mechanism behind the root-exudate driven microbial ecology. Our study uses rice as a model system to determine the importance of plant physiological traits (including root exudation rates and composition) on nitrification dynamics and BNI efficiency. This approach will enable to address important research gaps, such as those related to the spatial localisation of the BNI effect in soil, the physiological mechanism of the BNI efficiency in different plant genotypes or the range of microbes inhibited by BNI.

This study tested the following hypotheses: (H1) Secretion of root BNI compounds leads to greater inhibition of nitrifiers in the rhizosphere than in the bulk soil, with a stronger inhibition of AOA than AOB in the rhizosphere; (H2) The differential BNI efficiency of different rice genotypes is mediated by the rate and chemical composition of their root exudation; (H3) An AO culture bioassay including multiple AOA and AOB strains accurately models the variability in rice genotype BNI efficiency in soils; (H4) the BNI compound in high BNI rice exclusively inhibits AO.

## Results

### Ammonia oxidiser inhibition in the rice rhizosphere

AOA ammonia monooxygenase subunit A (*amoA*) gene abundance was nine times greater than AOB *amoA* gene abundance at the beginning of the incubation. AOA abundance remained higher than AOB abundance during the 90-days incubation (Fig. S1) (<u>Statistic A:</u> ANOVA; four-way interaction between AO group, rice genotypes, soil compartment and time; *F*-value= 4.1; *P*-value = 1.3×10^-3^). The planted/control ratio of AO abundance decreased significantly over time in the rhizosphere compared to bulk soil (Fig. 1, S2) (<u>Statistic B:</u> ANOVA; four-way interaction between AO group, rice genotypes, soil compartment and time; *F*-value= 209.8; *P*-value = 2.0×10^-16^). This difference between bulk and rhizosphere compartments suggests that AO are more inhibited in the rhizosphere than in the bulk soil. For both AOA and AOB analysed independently, the *amoA* AO abundance was not statistically different within (i.e., the rhizosphere soil compartment) and outside (i.e., the bulk soil compartment) the mesh bag in the control pots throughout the experiment (Fig. S3) (<u>Statistic C:</u> ANOVA; three-way interaction between AO group, soil compartment and time; *F*-value= 0.1; *P*-value = 0.96). This suggests that the mesh bag used in the experiment to separate the bulk and the rhizosphere soil compartments did not affect the motility of the microorganisms in the soil).

**Figure 1:**
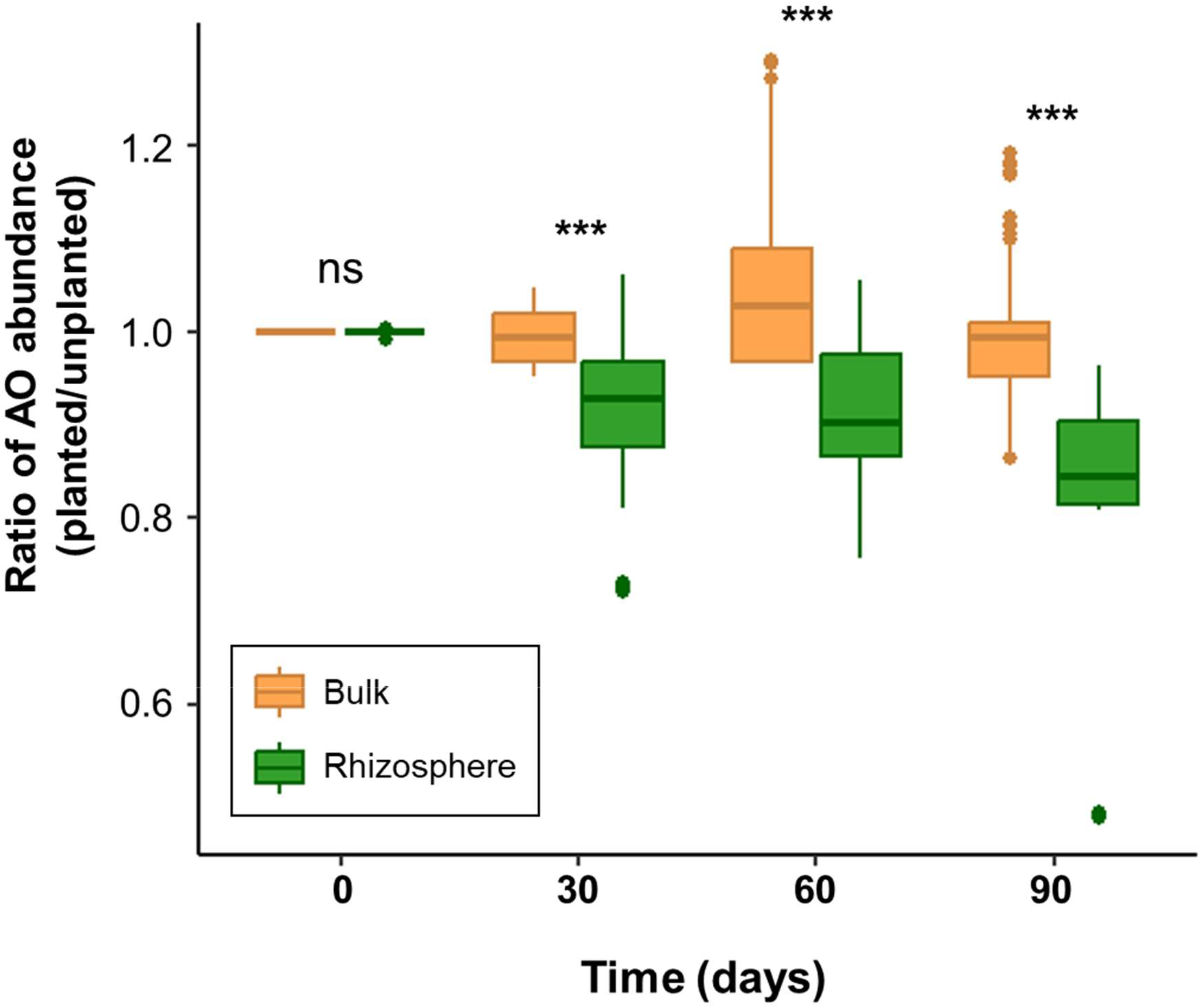
Ammonia oxidiser (AOA or AOB) ratio (planted/unplanted) of abundance in the bulk and rhizosphere soil compartments over time for the five rice genotypes. Data are represented as boxplots, and ‘ns’ and ‘****’ on top of each boxplot pair denote significant differences between bulk and rhizosphere ratio of abundance with *P*-value > 0.05 and *P*-value = 2×10^-16^, respectively, tested using Statistic B.

The AO inhibition was inferred from the decrease in the ratio of AO abundance (AOA or AOB) over time in the rhizosphere of each rice genotype, termed as death rate (Table S1). The death rate was statistically different between the two groups, AOA and AOB, and this effect depended on the rice genotype (Fig. 2A) (<u>Statistic D:</u> ANOVA; two-way interaction between AO group and rice genotype; *F*-value= 727.3; *P*-value = 2.0×10^-16^). The BNI efficiency for the five rice genotypes was then inferred using the sum of AOA and AOB death rates; this provided one high-BNI genotype (Lemont), three intermediate-BNI genotypes (IR64, Koshihikari and Moroberekan) and one low-BNI genotype (Azucena) (Fig. 2B) (<u>Statistic E:</u> ANOVA; rice genotype effect; *F*-value= 13981; *P*-value = 2.0×10^-16^).

**Figure 2:**
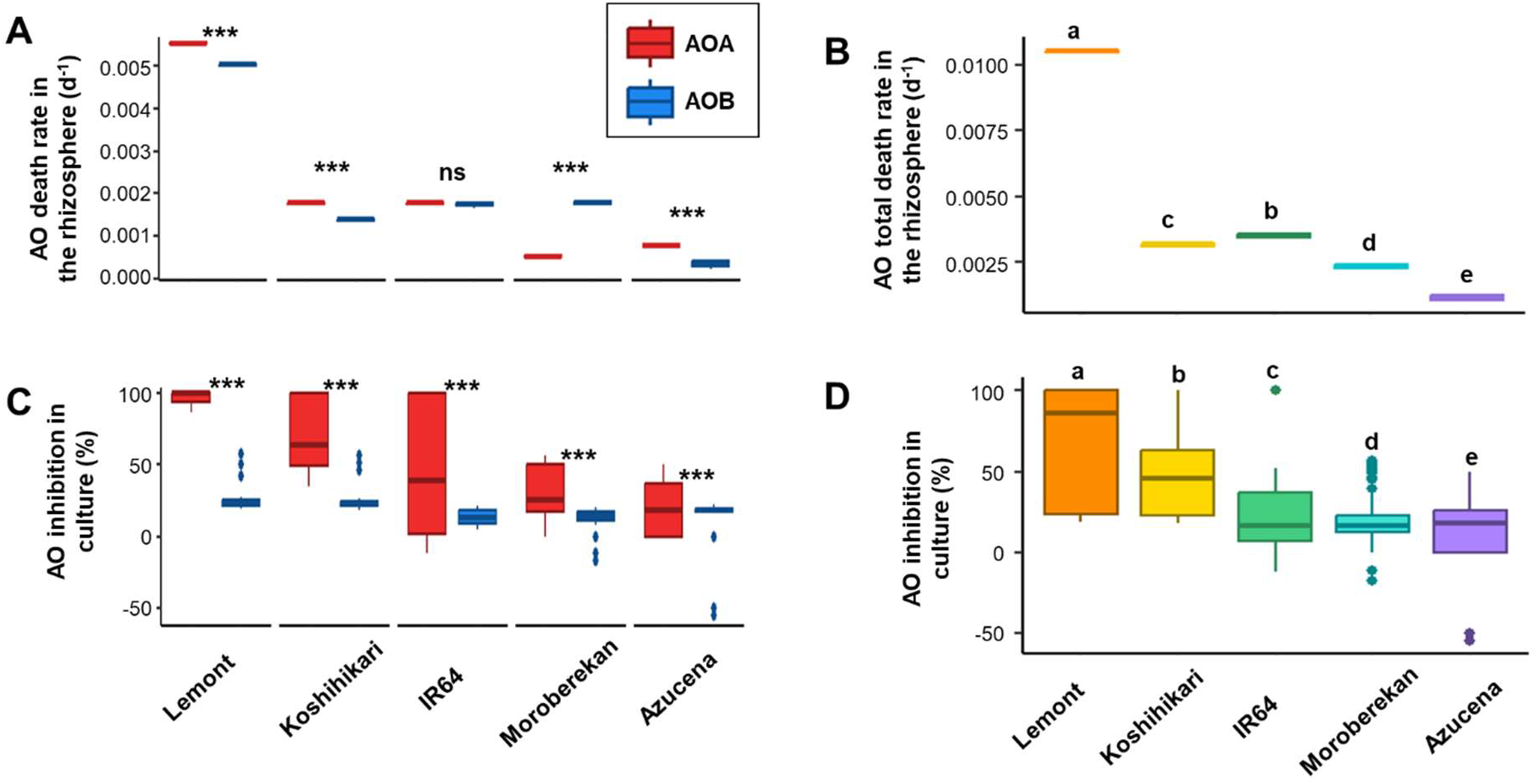
Estimation of Biological Nitrification Inhibition (BNI) efficiency in the five different rice genotypes using different approaches. (A) The AOA and AOB specific death rate was estimated over 90 days in the rhizosphere of the five rice genotypes. (B) The BNI efficiency in the rhizosphere of the five rice genotypes was estimated by the total AO (AOA + AOB) death rate. (C) The AOA and AOB growth rate inhibition was estimated in culture bioassay using root exudates from the five rice genotypes. (D) The BNI efficiency of the five rice genotypes was estimated by the growth rate inhibition of AO cultures by rice root exudates. For panels C and D, the AO strains were AOB – *N. europaea, N. multiformis* and *N. tenuis* and AOA – *N. viennensis*, *Ca.* N. sinensis and *Ca.* N. franklandus. For panels A and C, ‘ns’ and ‘****’ on top of each AOA and AOB box pair denote significant differences in BNI efficiency between AOA and AOB within each sub-plot, with *P*-value > 0.05 and *P*-value = 2×10^-16^, respectively, tested using Statistic D for panel A and Statistic J for panel C. For panels B and D, different letters on top of each box denote significant differences (*P*-value < 0.01) in BNI efficiency between the different rice genotypes, tested using Statistic E for panel B and Statistic J for panel D.

### Importance of root exudates for rice BNI efficiency and characterisation of a novel BNI compound

The five rice genotypes differed physiologically, particularly for their dry root weight (<u>Statistic F:</u> ANOVA; rice genotype effect; *F*-value= 173.5; *P*-value = 3.4×10^-9^). Similarly, the root exudate biomass also varied between rice genotypes (<u>Statistic G:</u> ANOVA; rice genotype effect; *F*-value= 4582; *P*-value = 2.2×10^-16^). The root exudate biomass is negatively correlated with the dry root weight (Fig. 3A) (<u>Statistic H:</u> linear model; r^2^=0.54; *F*-value= 17.1; *P*-value = 1.0×10^-3^), suggesting that the plants producing more root exudates have a lower root mass. These physiological traits were incorporated into a single index, the exudation rate, representing the mass of root exudates secreted by the plant root system over time. The root exudate rate was positively correlated with the total AO death rate in the rhizosphere (Fig. 3B) (<u>Statistic I:</u> linear model; r^2^=0.93; *F*-value= 177.1; *P*-value = 6.0×10^-9^), suggesting that plants with a higher root exudation rate have a higher BNI efficiency.

**Figure 3:**
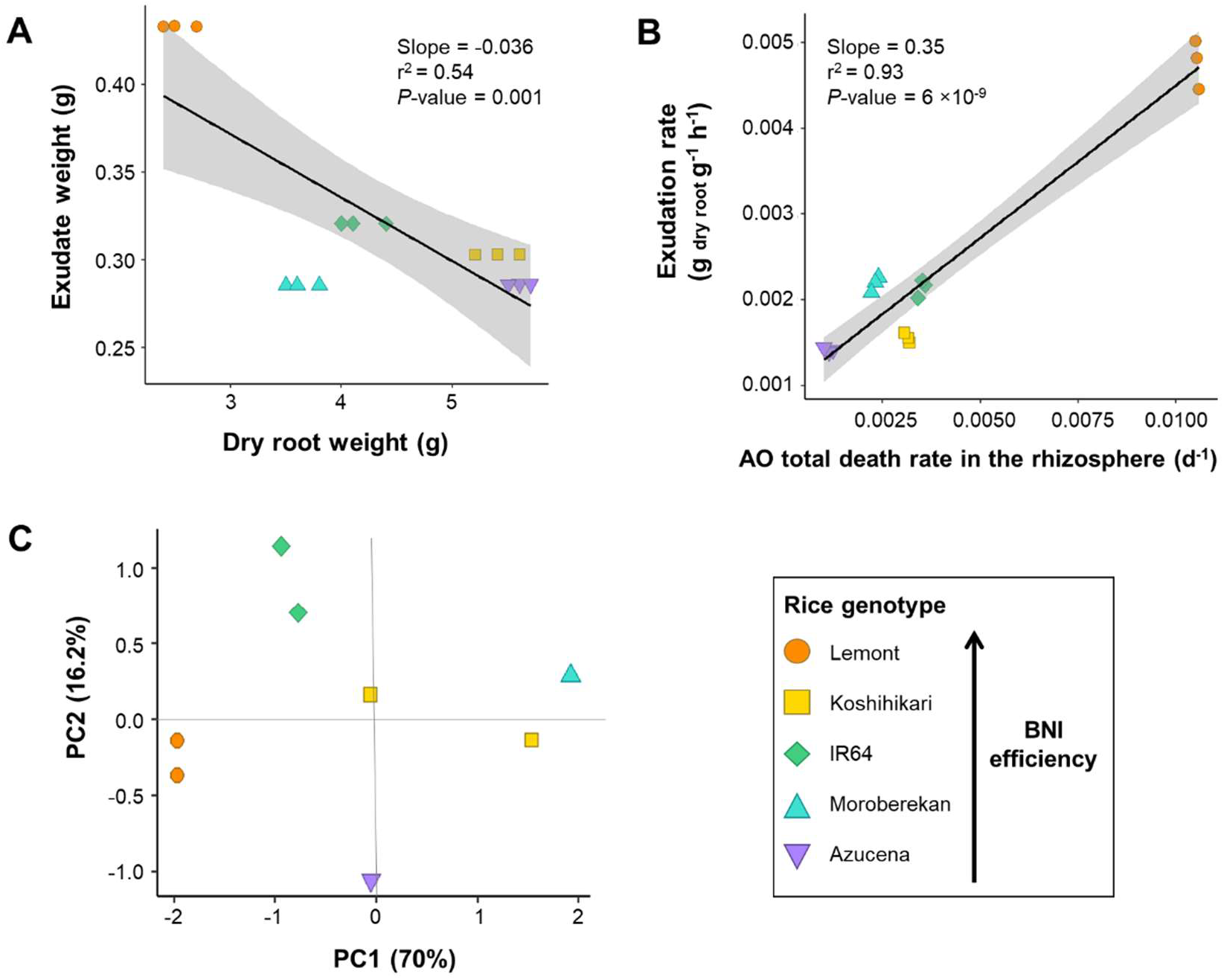
Influence of plant physiology on the root exudate rate and composition. Panel A shows the negative correlation between dry root weight and root exudate weight for the five rice genotypes. Panel B shows the positive correlation between root exudation rate and total AO death rate of the five rice genotypes. On panel C, the Principal Component Analysis (PCA), based on the LC-MS root exudate composition, represents the diversity of 90-days excreted compounds for the five rice genotypes. Each point indicates a plant replicate and 3 replicates per genotype were included.

The 90-day root exudate LC-MS profile of the five rice varieties differed between the rice genotypes, with 86.2% of the variance between root exudate composition being attributed to differences between rice genotypes (Fig. 3C). The LC-MS profile of the rice genotypes indicated a peak at 3 min in the positive ionisation profile, having a mass-to-charge ratio of 242.2838 corresponding to the protonated molecular ion [M^+^H]^+^ with formula C_16_H_36_N (Fig. 4). *N*-butyldodecane-1-amine was identified based on the Reaxys database and verified by comparing the tandem mass spectrometry (MS-MS) fragmentation pattern with the published data for *N*-butyldodecane-1-amine (Wei et al., 2018) as well as with the MS-MS data of the purchased pure compound (Fig. 4, S5).

**Figure 4:**
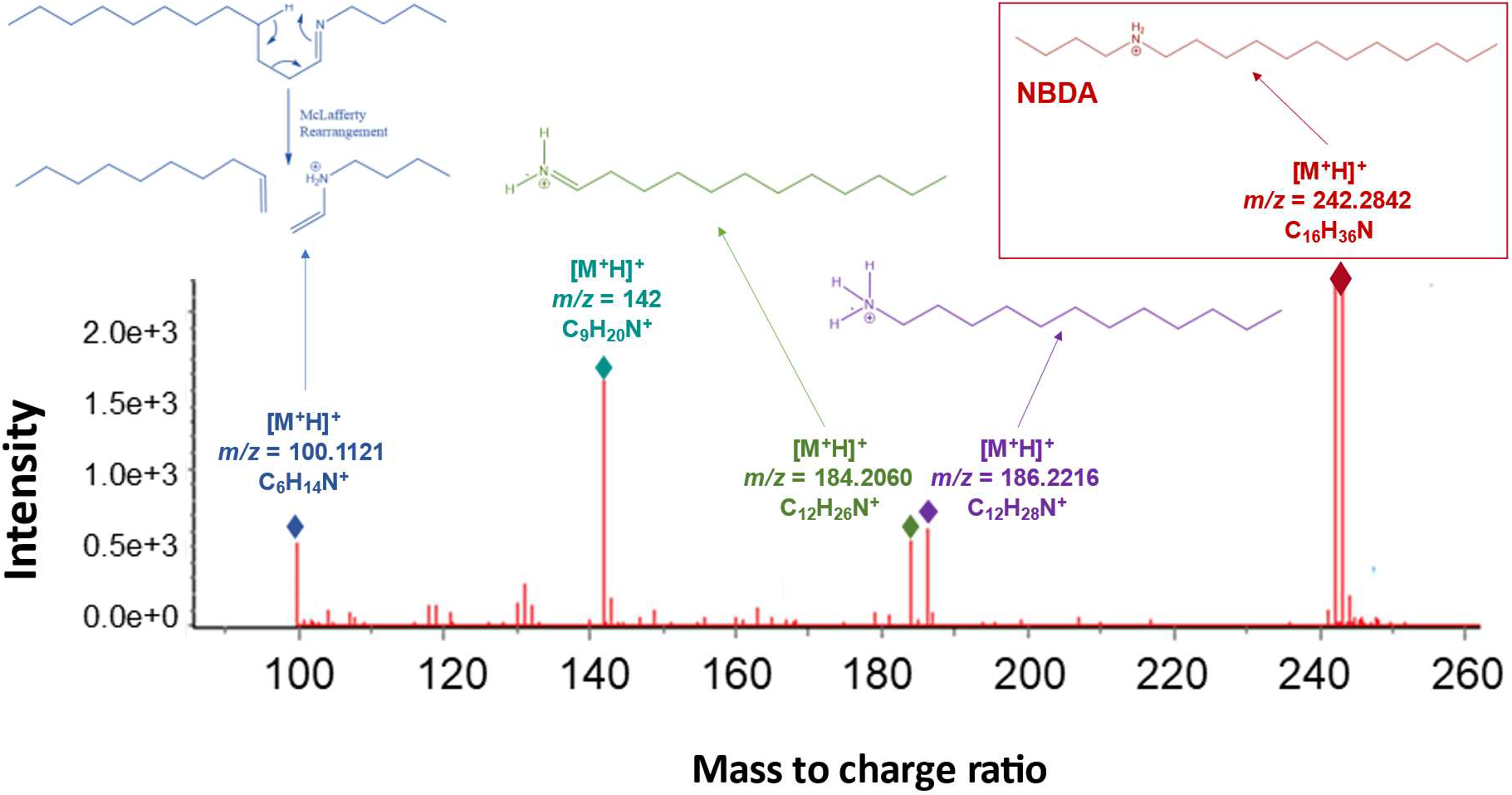
LC-MS guided isolation of *N*-butyldodecan-1-amine (NBDA) LC-MS chromatogram region and fragmentation pattern of the compound NBDA obtained in the positive ESI-MS mode (UV detection at 228 nm) of the Lemont root exudates. The exact mass and structure of fragments observed in the LC-MS chromatogram are indicated above each fragment peak.

### Modelling of the rice genotype BNI efficiency using an AO culture bioassay

The BNI efficiency of the different rice genotypes was evaluated by using AOA and AOB culture inhibition bioassays, as previously validated (Kaur-Bhambra et al., 2022). The BNI efficiency of the different rice genotypes was significantly different between the different rice genotypes (Fig. 2D) (<u>Statistic J</u>, ANOVA, rice genotype effect; *F*-value=739.1; *P*-value = 2.0×10^-16^), and this effect depended on the AO groups and the strains considered (Fig. 2C, S4) (<u>Statistic J</u>, ANOVA, three-way interaction between AO group, strain and rice genotype; *F*-value= 170.7; *P*-value = 2 .0×10^-16^). The different rice genotypes showed a similar ranking of BNI efficiency in the bioassay than using AO total death rate in the rhizosphere (Fig. 2B, D).

### BNI efficiency of the novel rice BNI compound, *N*-butyldodecane-1-amine

The BNI efficiency of NBDA was tested against AOA and AOB cultures using the pure compound (Fig. 5A, S3). NBDA has a very high BNI efficiency (range 1.3 - 49.8 μM for the six strains tested). AOA strains were significantly more inhibited by NBDA than AOB strains (Fig. 5A) (<u>Statistic K</u>, ANOVA, strain effect; *F*-value= 872; *P*-value = 2.0×10^-16^). NBDA had a 30-times greater BNI efficiency than another rice BNI, 1,9-decanediol (Kaur-Bhambra et al., 2022).

**Figure 5:**
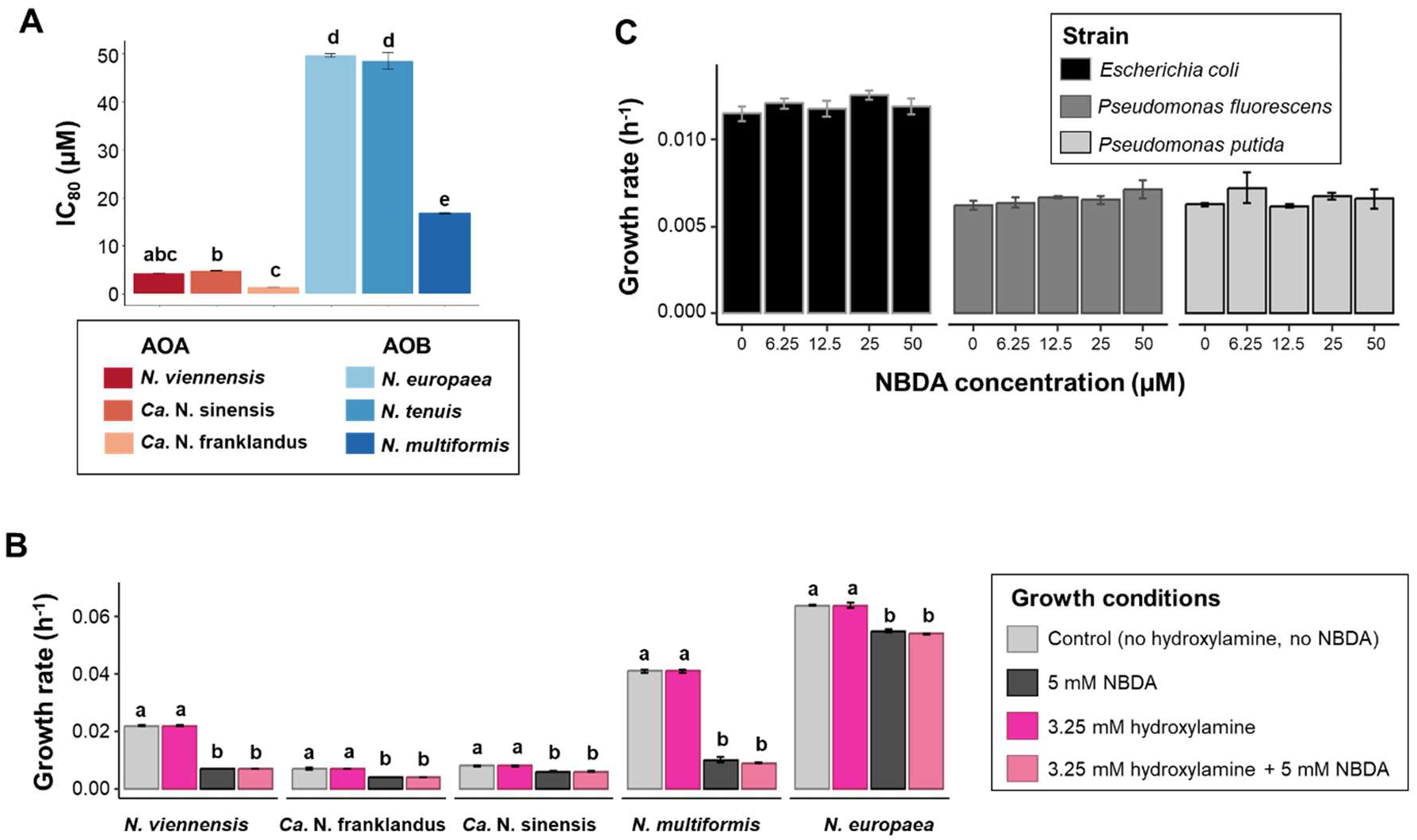
Inhibition potential of rice BNI, *N*-butyldodecan-1-amine (NBDA) (A) Inhibitory concentrations at 80% inhibition (IC_80_) of six ammonia-oxidiser strains (3 AOA and 3 AOB) for NBDA. (B) Comparison of the growth rates of five ammonia oxidising strains with and without hydroxylamine supplementation in the presence and absence of NBDA. AO strains used in the study are, AOB – *N. europaea, N. multiformis* and *N. tenuis* and AOA – *N. viennensis*, *Ca.* N. sinensis and *Ca.* N. franklandus. (C) Comparison of growth rates of three non-AO strains, *Escherichia coli* MG1655, *Pseudomonas fluorescens* NCIMB 10586 and *Pseudomonas putida* PG NCIMB 9816, in the presence of different concentrations of NBDA. All data are presented as bar plots, standard errors (n=4) are indicated by error bars, and different letters within each subplot denote significant differences (*P*-value < 0.05) in inhibition based on Statistic K, L and M, for panel A, panel B and panel C, respectively.

The mechanism of NBDA ammonia oxidiser inhibition was investigated by focusing on the hydroxylamine oxidoreductase (HAO) in AOB and an equivalent enzyme(s) in AOA (currently unidentified), with hydroxylamine being the intermediate compound in both groups of ammonia oxidisers. The inhibition by NBDA was not alleviated by the addition of hydroxylamine in the AO strains tested (Fig. 5B) (<u>Statistic L</u>, LM contrast medium (NBDA) – medium (hydroxylamine + NBDA); *P*-value = 0.37).

The inhibition efficiency of NBDA was tested on three non-AO strains, *E. coli* MG1655*, P. putida* NCIMB 9816 and *P. fluorescens* NCIMB 10586. NBDA had no inhibitory effect on the growth rate of these strains, at least up to a concentration of 50 µM (Fig. 5C) (<u>Statistic M</u>, ANOVA, Two-way interaction between strain and NBDA concentration; *F*-value= 1.0; *P*-value = 0.44).

## Discussion

### Greater inhibition of nitrifiers in the rhizosphere than in the bulk soil

As BNIs secreted as root exudates are primarily exudated in the rhizosphere, we hypothesised that the nitrifier inhibition is greater in the rhizosphere than in the bulk soil (H1). We confirmed this hypothesis (Fig. 1), contradicting previous findings in rice paddy systems (Ke et al., 2013; Li et al., 2007). In our bulk soil sampling, we homogenised the top and bottom soil layers, hereby including surface soils where AOA abundance is generally greater due to oxygen availability (Ke et al., 2013). Such discrepancy in inhibition between the rhizosphere and bulk soils was previously observed within other BNI-producing crops, such as sorghum (Watanabe et al., 2015; Burnham et al., 2022) and ryegrass (Simon et al., 2020), providing a stronger framework of BNI mechanism in soil. This high BNI efficiency in the rhizosphere has been previously linked to rhizosphere acidification during active ammonium uptake and BNI release (Watanabe et al., 2015; Di et al., 2018; Afzal et al., 2020). However, the BNI release of 3-epi-brachialactone was shown to be directly regulated by the generated proton motive force defining transport rates across the plasma membrane rather than indirectly linked to pH and ammonium availability in *Brachiara humidicola* (Egenolf et al., 2021). Other mechanisms related to AO niche specialisation likely favour the different abundance of AO between rhizosphere and bulk soils, as the rhizosphere environment encounters recurrent pH fluctuating conditions in soil microniches (Watanabe et al., 2015).

### Stronger inhibition of AOA than AOB in culture bioassay but not in the rhizosphere

Similarly to numerous rice acidic paddy soils (Chen et al., 2008; Ke et al., 2013; Hu et al., 2015), AOA were the more abundant nitrifiers in our study. The greater adaptation of AOA than AOB to thrive under low pH, lower ammonium availability and low oxygen concentration environments are likely the key factors influencing their dominance under hypoxic environments (Kraft et al., 2022; Xie et al., 2021; Martens-Habbena et al., 2009). AOA dominance does not preclude AOB in this type of soils (Chen et al., 2011; Ke et al., 2013; Huang et al., 2014; Zhang et al., 2019; Fu et al., 2020). We hypothesised that the inhibition of AOA will be greater AOB in the rhizosphere (H1) (Kaur-Bhambra et al., 2022), and this was confirmed in culture bioassays (Fig. 2C). While confirmed in soil for some rice genotypes, AOA did not appear to be preferentially inhibited compared to AOB in all tested plant genotypes (Fig. 2A). Explaining the mechanism behind such soil *vs* culture bioassay discrepancy is difficult to advance but could be due to specific genotype exudates favouring different AO members. The restricted range of AO cultures available prevents direct testing in cultures. We could confirm that our current AO culture bioassay, including multiple AOA and AOB strains models, nicely the variability in rice genotype BNI efficiency in soils (H3) (Fig. 2B, D), but a greater diversity of AO strain would likely increase the accuracy of this approach. In addition, the respective AOA and AOB community structure in the soil rhizosphere between high- and low-BNI rice genotypes should be further investigated in our current study.

The BNI efficiency retrieved in the five rice cultivars tested in our study differed from a previous study (Tanaka et al., 2010). Multiple factors can explain this discrepancy. First, the estimation of BNI efficiency were performed using different approaches: bioluminescent *N. europaea* inhibition by root exudates in Tanaka et al. (2010) *versus* multiple AOA and AOB inhibition by root exudates and AO inhibition in soil in our study. The two studies also used different plant growth systems (soil versus sterile hydroponic). The sterile hydroponic conditions alter the root physiology and negate all plant-microbial interactions, involved in the root exudate composition (Ascough & Fennell, 2004; Sasse et al., 2020; Tavakkoli et al., 2010), leading to a distinct root exudates than in soil. Finally, root exudates were collected at different plant stages in the two studies (90 *versus* 60 days), while plant developmental stage influences oxygen leakage and root exudation (Ikenaga et al., 2003; Ke et al., 2013; Tanaka et al., 2010).

### Influence of plant physiology on BNI efficiency

A wide range of BNI efficiency was demonstrated among different plant cultivars within the same species, for rice (Chen et al., 2022; Illarze et al., 2021; Li et al., 2007; Tanaka et al., 2010; Sun et al., 2016; Zhang et al., 2022; this study) and in other crops, such as pearlmillet (Ghatak et al., 2022), maize (Mwafulirwa et al., 2021), koronivia grass (Vázquez et al., 2020) and sorghum (Gao et al., 2022). We hypothesised that the differential BNI efficiency of different rice genotypes is mediated by their root exudation rate and chemical composition (H2). Variable root exudates were previously observed between different genotypes of wheat (Iannucci et al., 2021; Nguyen et al., 2019), koronivia grass (Nakamura et al., 2020), sorghum (Seitz et al., 2022) and pearl millet (Ghatak et al., 2022), and a suggested link between BNI efficiency and root exudation was proposed in drought-affected millet (Ghatak et al., 2022). In our study, the different rice genotypes used showed different root exudation masses, themselves negatively correlated to root biomass (Fig. 3A). Such negative correlation was also observed in members of the *Pooideae* subfamily (grasses and forbs) (Guyonnet et al., 2018). This suggests that C3 plant physiological traits might control for the biomass of root exudates secreted. In addition, the chemical composition of the root exudates differed between the five tested rice genotypes (Fig. 3C). We confirmed that the root exudate rate and chemical composition can therefore, at least partially, explain the distinct BNI efficiencies in the different rice genotypes (Fig. 3B, C). Plant life strategy in those C3 plants could explain these genotype-specific BNI efficiencies, and this needs to be further investigated. In addition, as root systems are dynamic in nature, possible changes in root exudation mass and chemical composition would change depending on the plant growth stages (Gransee & Wittenmayer, 2000; Tanaka et al., 2010), growth conditions (Ghatak et al., 2022; Maurer et al., 2021) and associated microbial community (Eisenhauer et al., 2017), so a greater range of conditions should be analysed.

### Nitrification inhibitory activity of the novel BNI compound, *N*-butyldodecane-1-amine

The identified BNI compound N-butyldodecane-1-amine (NBDA), is a linear aliphatic amine, representing a new class of BNI compounds isolated from plants. NBDA is structurally similar to previously isolated fatty acid BNIs, such as 1,9 decanediol isolated from rice roots (Sun et al., 2016) or linoleic acid and linolenic acid extracted from shoot tissues of *Brachiaria humidicola* (Subbarao et al., 2008). NBDA shows relatively higher inhibitory activity than other known rice BNI compounds (Kaur-Bhambra et al., 2022).

NBDA did not inhibit non-AO bacteria, suggesting the compound activity may be restricted to ammonia oxidisers, through direct enzymatic ammonia oxidation inhibition. Our results suggest that NBDA may inhibit the second and/or the third step of the ammonia oxidation pathway, which mediates the conversion of hydroxylamine to nitrite via nitric oxide (Caranto & Lancaster, 2017). Nonetheless, the precise mode of action of most BNI compounds, including NBDA, still remains to be elucidated. One of the important future challenge for the BNI field would be to determine the proteins and enzyme kinetics involved in BNI.

## Materials and methods

### Soil sampling and experimental setup

Sandy loam agricultural soil of pH 5.5 was sampled on 28 April 2021 from the upper 10 cm of a long-term pH-maintained plot in SRUC, Craibstone, Scotland (grid reference NJ872104). Soil characteristics as described previously (Kemp et al., 1992). The soil was dried overnight at 40°C, sieved with a 3.35 mm mesh size and stored at 4°C before the construction of microcosms. Microcosms were prepared in black polyethylene containers (height, 22 cm; diameter, 17 cm) with 1.5 kg of dry soil per microcosm. Nylon mesh bags (2.5 mm mesh, 18 cm height; 15 cm diameter) (obtained from Normesh, UK) containing 0.5 kg of dry soil were placed into the centre of each pot containing 1 kg of soil, creating two independent soil compartments through which water, nutrients and microorganisms could pass freely. The plants were placed in the bags; hence, the central-rooted and the outside non-rooted compartments contained the rhizosphere and the bulk soils, respectively. The microcosms were set up in a greenhouse with 65% humidity, a temperature regime of 35°C: 28°C, light: dark photoperiod of 14 h: 10 h and light intensity of 400 μmol m^−2^ s^−1^ provided by high-pressure sodium lamps. After 2 days of water-logging, KCl, Na_2_HPO_4_ and NH_4_Cl were applied as basal fertilisers at 17 mg K and 50 mg P and 100 mg N per kg of dry soil.

The rice varieties Lemont, IR64, Koshihikari, Azucena and Moroberekan were chosen to represent a spectrum of BNI efficiency based on a previous study (Tanaka et al., 2010). Seeds of rice varieties were sterilised with 5% H_2_O_2_ for 15 min, rinsed and soaked with deionised water for 2 h. The seeds were germinated on filter paper in Petri dishes at 40°C in the dark. After 4 days of incubation, the seeds were placed under light till shoots reached a length of about 1 cm. Two days after fertilisation, the germinated rice seeds were transplanted into the pots with three seedlings per pot. The seedlings of the selected rice varieties were grown for 90 days in soil microcosms where a nylon mesh bag separated the bulk and rhizosphere soil compartments. A set of microcosm pots were incubated without rice plants as control. Each treatment (five rice varieties and one control) contained triplicate pots for the four time points (at 0, 30, 60 and 90 days).

### Molecular analyses

Triplicate treatments of bulk soil, rhizosphere soil and plants were destructively sampled at each time point. The nylon mesh bag was removed from the pot, and the soil inside the nylon bag, closely adhering to the roots, was sampled as rhizosphere soil. The upper and lower 3 cm soil layers outside the mesh bag were collected, mixed, and designated as bulk soil. Soils were stored at - 80°C for molecular analyses. DNA was extracted from 0.5 g soil using a phenol-chloroform extraction method (Griffiths et al., 2000; Nicol et al., 2008). Extracted DNA was cleaned with AMPure® XP magnetic beads (Invitrogen, USA), according to the manufacturer’s instructions. DNA concentrations and quality were measured using a Nanodrop (ND-1000 Spectrophotometer, Labtech). qPCR was used to determine the abundance of archaeal and bacterial *amoA* gene using crenamoA23F/crenamoA616R (Tourna et al., 2008) and amoA-1F/amoA-2R (Rotthauwe et al., 1997) primer pairs, respectively, using standards and cycling conditions as described previously (Thion & Prosser, 2014). Amplification efficiencies for archaeal and bacterial *amoA* qPCR assays were 87% and 96%, respectively, with r^2^ values > 0.99. The *amoA* gene abundance were adjusted for dry soil weight and converted to their natural log values for further analyses.

The ratio of the AO abundance in the rhizosphere and bulk compartments were assessed separately for AOA and AOB at each time point, using the log converted AO abundance in each condition (rice genotype and soil compartment) relative to the AO abundance in control in the corresponding soil compartment. For each rice genotype, the BNI efficiency in soil was determined by applying a linearly model of the decrease in AO abundance (of AOA and AOB separately) over time in the rhizosphere soil compartment. The absolute value of the model slope represented the death rate of the corresponding AO, and the sum of the AOA and AOB death rate corresponded to the total AO death rate in the rhizosphere. Significant differences in (a) the *amoA* gene abundance between two AO groups, (b) the changes in the ratio of AO abundance between the bulk and the rhizosphere soil compartments, and (c) the AO death rate between the two AO groups for the different rice genotypes were estimated using multi-factorial linear modelling and two-way ANOVA tests, followed by post hoc tests using emmeans (Russell, 2022) in R V4.1.2.

### Root exudate collection

The root exudates were collected from 90-day-old rice plants using triplicate microcosms per genotype. The plants were submerged in water for 1 h to loosen the soil particles around the root. The roots were then rinsed thoroughly with deionised water to remove most of the soil particles attached to the plants. The plants were then gently placed in a clean collection solution comprising 0.1 M CaCl_2_ and 0.01 M NH_4_Cl in deionised water for 5 h to recover from manipulation stress (Williams et al., 2021). The plants were transferred into dark flasks containing 1 L of collection solution for 36 h. During this period, the collection solution was repeatedly replenished and maintained at a volume of 1 L. After 36 h, the collection solutions were filtered using a 0.22 μm filter and stored at 4°C for no more than 6 days until further treatment. The filtered samples were evaporated until dry using a rotary evaporator (Buchi Rotary Evaporator R-114 Series) at 40°C. The residues remaining in the round-bottom flask were dissolved in 10 mL of 50% HPLC grade methanol and re-evaporated to dryness under N_2_. The collected dry root exudate mass was weighed and stored at −20 °C.

To reduce the bias of leftover soil particles attached to plant roots in BNI compound and activity determination, 0.5 g of soil from control pots was mixed with 1 L of collection solution and processed as a control. Plant root length, root width and shoot length were recorded at the end of root exudate collection. The roots and shoots were washed, separated, and dried in an air oven at 65°C for 24 h to determine the plant’s dry root and shoot weight. Significant differences in (a) the dry root weight and (b) the root exudate mass between the different rice genotypes were measured using a one-way ANOVA. Simple linear modelling was used to analyse the correlation and the significance of the interactions between (a) the dry root weight and the root exudate mass and (b) the root exudate rate and the AO death rate in soil for the different rice genotypes.

### Identification of BNI compounds

Chemical characterisation of the root exudates was performed by LC-MS analysis, using 1 mg of the dried root extracts re-dissolved into 50% HPLC grade methanol. LC-MS analysis was carried out using an Agilent 6490 QqQ equipped with a High-Performance Liquid Chromatography (HPLC) (Agilent 6490) system equipped with a reversed-phase C18 analytical column and eluted with a mobile phase starting with 5% acetonitrile (containing 0.1% formic acid) followed by a gradient up to 100% acetonitrile (containing 0.1% formic acid) for 15 min, at a flow rate of 1 mL/min. The subsequent eluate was passed into a mass spectrometer with a Bruker maXis II Quadrupole-Time-of-Flight (Q-TOF) mass analyser. Mass spectra were measured in positive ion mode electrospray ionisation of triplicate root exudate for each genotype at day 90.

Compounds were identified based on their retention time using Bruker Daltonics Data Analysis software. Mass spectra dereplication was performed using the Reaxys database to match observed fragment masses with literature values. Peaks of interest were selected as those present in all rice genotypes. Among those, a compound was identified as a new BNI compound, and its molecular formula was verified by comparing the retention time with an authentic commercially produced compound. To understand changes in the root exudation, the triplicate MS spectra for the different genotypes were centroided using msConvert (Adusumilli & Mallick, 2017) and Mzmine3 (Pluskal et al., 2010) and the peak areas were compared using K-nearest Neighbours algorithm principal coordinate analysis in the MetaboAnalyst5 (Xia & Wishart, 2011) (see details in Supplementary methods).

### Determination of the inhibitory effect of root exudates and isolated compound NBDA

Three pure AOA and three AOB strains were selected to test the AO inhibitory activity of both the root exudates and the novel identified chemical BNI compound. The selection included the AOA strains *Nitrososphaera viennensis* (EN76), isolated from an Austrian pH 8 garden soil (Tourna et al., 2011), *Candidatus* Nitrosotalea sinensis (Nd2), isolated from a Chinese pH 4.7 paddy soil (Lehtovirta-Morley et al., 2014), and *Candidatus* Nitrosocosmicus franklandus (Nfr-C13), isolated from a Scottish pH 7.5 sandy loam soil (Lehtovirta-Morley et al., 2016);, and the AOB strains, *Nitrosospira tenuis* NV12, isolated from Hawaiian soil (Harms et al., 1976), *Nitrosomonas europaea* ATCC 19718 and *Nitrosospira multiformis* ATCC 25196, obtained from NCIMB (http://www.ncimb.com/). The three AOA strains were grown in their respective modified freshwater medium as described previously for *N. viennensis, Ca.* N. franklandus and *Ca.* N. sinensis (Lehtovirta-Morley et al., 2014, 2016; Tourna et al., 2011), at 35 °C, 40 °C and 35 °C, respectively. All three AOB strains, *Nitrosospira tenuis* NV12, *Nitrosomonas europaea* ATCC 19718 and *Nitrosospira multiformis* ATCC 25196 were grown using the modified Skinner and Walker medium (Skinner & Walker, 1961) at 28 °C.

For each crude root exudate, 0.1 g of exudates was initially dissolved in 1 ml of 50% filtered dimethyl sulfoxide (DMSO), and 10 µl of the dissolved root exudates were added to 5 ml of culture media to obtain a final concentration of 0.1% DMSO in culture media. This concentration was chosen to reduce any inhibitory effect caused by DMSO on the tested AO strains (Kaur-Bhambra et al., 2022). To remove any inhibition bias caused by trap solution salts or leftover soil phenolics, the dried control trap solution was also dissolved in 50% filtered DMSO and processed similarly to the other root exudates. The identified BNI compound, *N*-butyldodecane-1-amine (NBDA), was obtained from Sigma-Aldrich (purity of 96%). NBDA stock solutions were prepared using filtered 100% DMSO, and 5 µl of stock solution was added to a 5 ml medium at concentrations in the range of 0.2–50 μM, as required. All the treatments using root extracts and the pure compound were carried out in quadruplicates for each AO strain.

The ammonia oxidation enzyme inhibition by NBDA was analysed by incubating the AOA and AOB strains in the presence or absence of 3.25 mM hydroxylamine. It was assumed that the inhibitory effect would be alleviated in the presence of hydroxylamine if the BNI compound specifically inhibits the hydroxylamine oxidoreductase (HAO) enzyme or its equivalent in AOA (Subbarao et al., 2006). In addition, the inhibitory effect of NBDA was tested on three on-ammonia oxidisers strains: *Escherichia coli* MG1655, obtained from a stool sample of a diphtheria patient, *Pseudomonas fluorescens* NCIMB 10586, isolated from a UK soil, and *Pseudomonas putida* PG NCIMB 9816, isolated from a UK garden soil. The three non-AO strains were grown in LB growth media either at 35 °C for *E. coli* or at 25 °C for *Pseudomonas*.

Growth analyses and IC_80_ determination were performed as previously described (Kaur-Bhambra et al., 2022). Significant differences in (a) the inhibition between the different AO strains by root exudates; (b) the inhibition between AOA and AOB by root exudates; (c) the IC_80_ of NBDA between the different AO strains; (d) the growth rates with NBDA supplemented in the presence and absence of hydroxylamine and (e) the growth rates of non-AO strains in the presence of NBDA were measured using multi-factorial linear modelling and a two-way ANOVA followed by post hoc tests using emmeans (Russell, 2022) in R V4.1.2.

## Supplementary Figures

**Figure S1:**
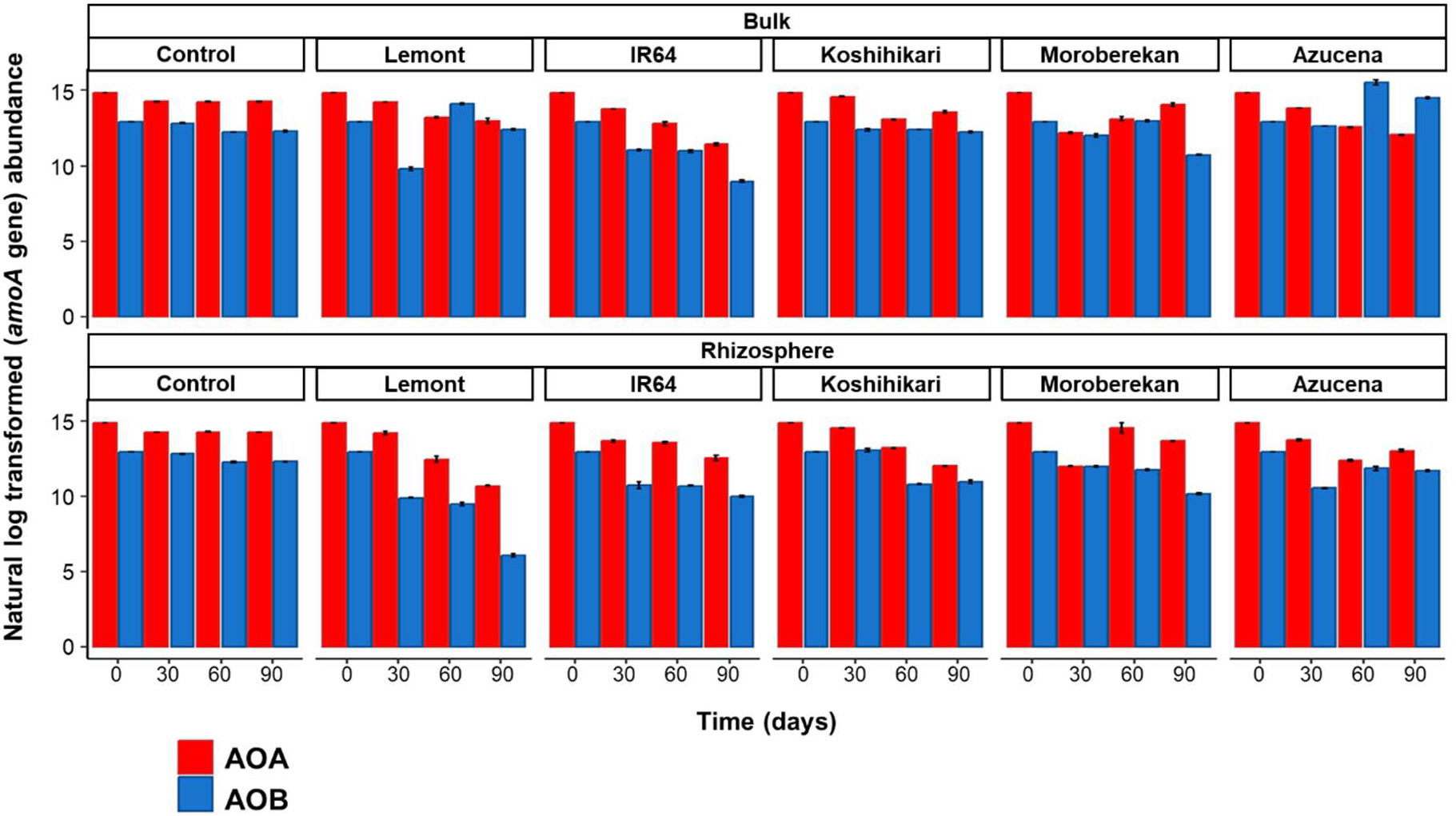
Abundance of ammonia-oxidisers in soil. The natural log-transformed AO abundances (AOA or AOB) in bulk and rhizosphere soil compartments over time in the five rice genotypes and control as determined by qPCR. Standard errors (n=3) are indicated by error bars.

**Figure S2:**
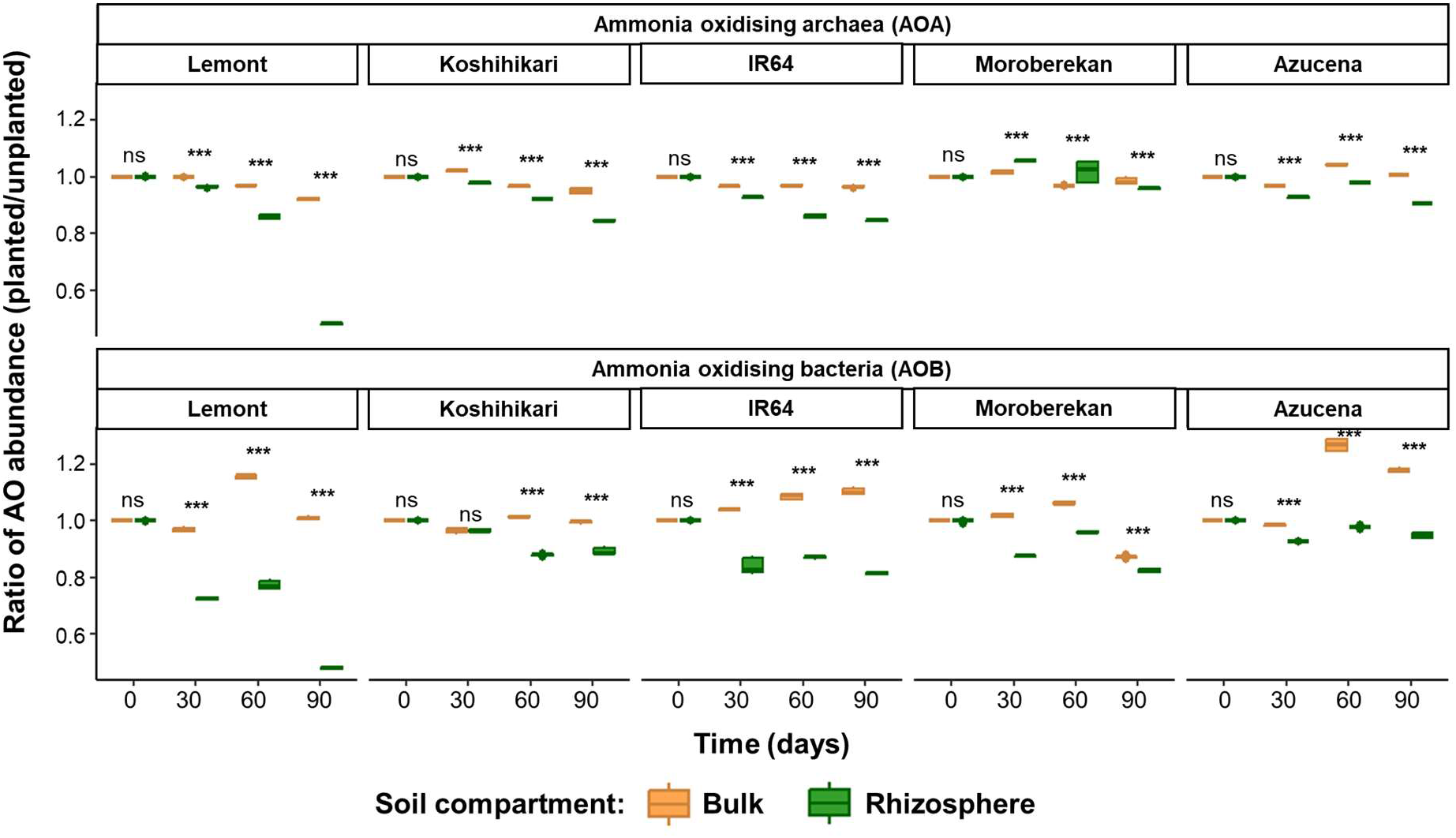
The ratio of abundance of ammonia oxidisers in the bulk and rhizosphere soil compartments. The ratio of ammonia-oxidiser (AOA and AOB) abundance in bulk and rhizosphere soil compartments over time for the five rice genotypes investigated. Data represented as a boxplot, ‘ns’ and ‘****’ on top of each boxplot pair denote significant differences in bulk vs rhizosphere ammonia-oxidiser abundance with *P*-value > 0.05 and *P*-value = 2×10^-16^, respectively, tested using Statistic B.

**Figure S3:**
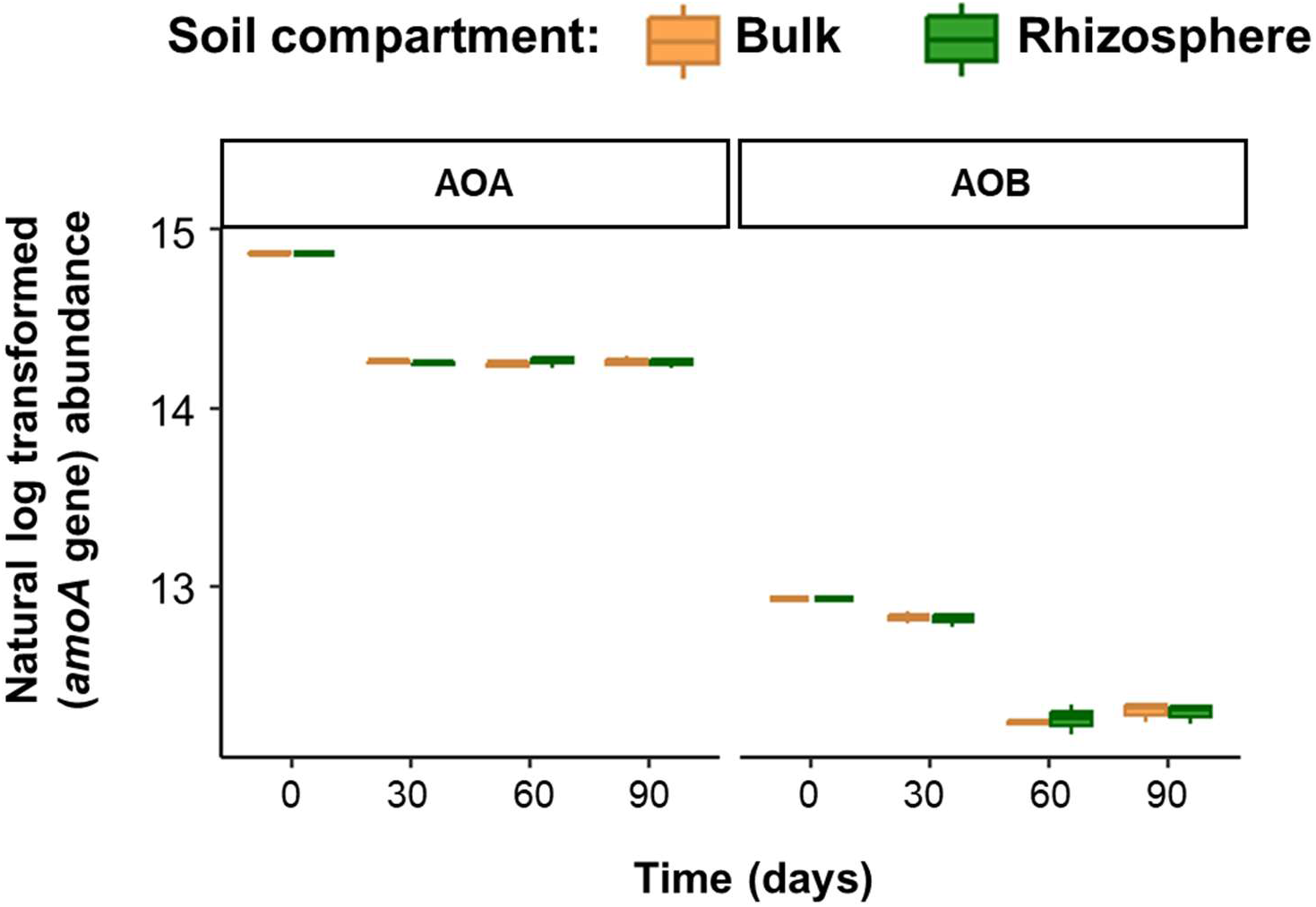
The abundance of ammonia oxidisers in the in the control soil microcosms. The log transformed ammonia-oxidiser (AOA and AOB) abundance in the bulk and rhizosphere soil compartments over time in the control pots.

**Figure S4:**
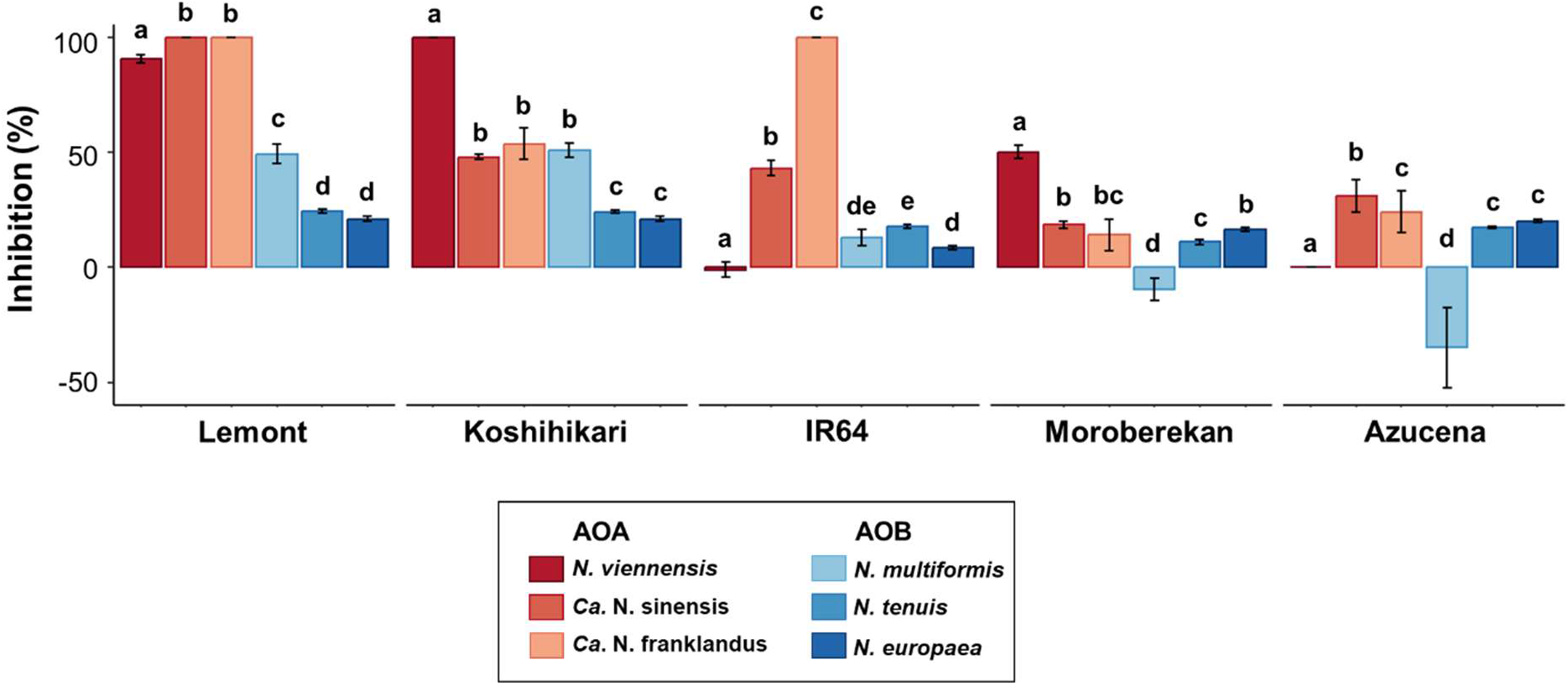
Biological Nitrification Inhibition of ammonia oxidisers in culture by rice root exudates. Inhibition of ammonia-oxidisers (AOA or AOB) in culture using an equal concentration of root exudates from the five rice genotypes inverstigated. Data presented as barplot and standard errors (n=16) are indicated by error bars. Different letters on top of each box denote significant differences (*P*-value < 0.01) in the AO inhibtion between the different rice genotypes, tested using Statistic J. Strains used in the study are, AOB – *N. europaea, N. multiformis* and *N. tenuis* and AOA – *N. viennensis*, *Ca.* N. sinensis and *Ca.* N. franklandus.

**Figure S5:**
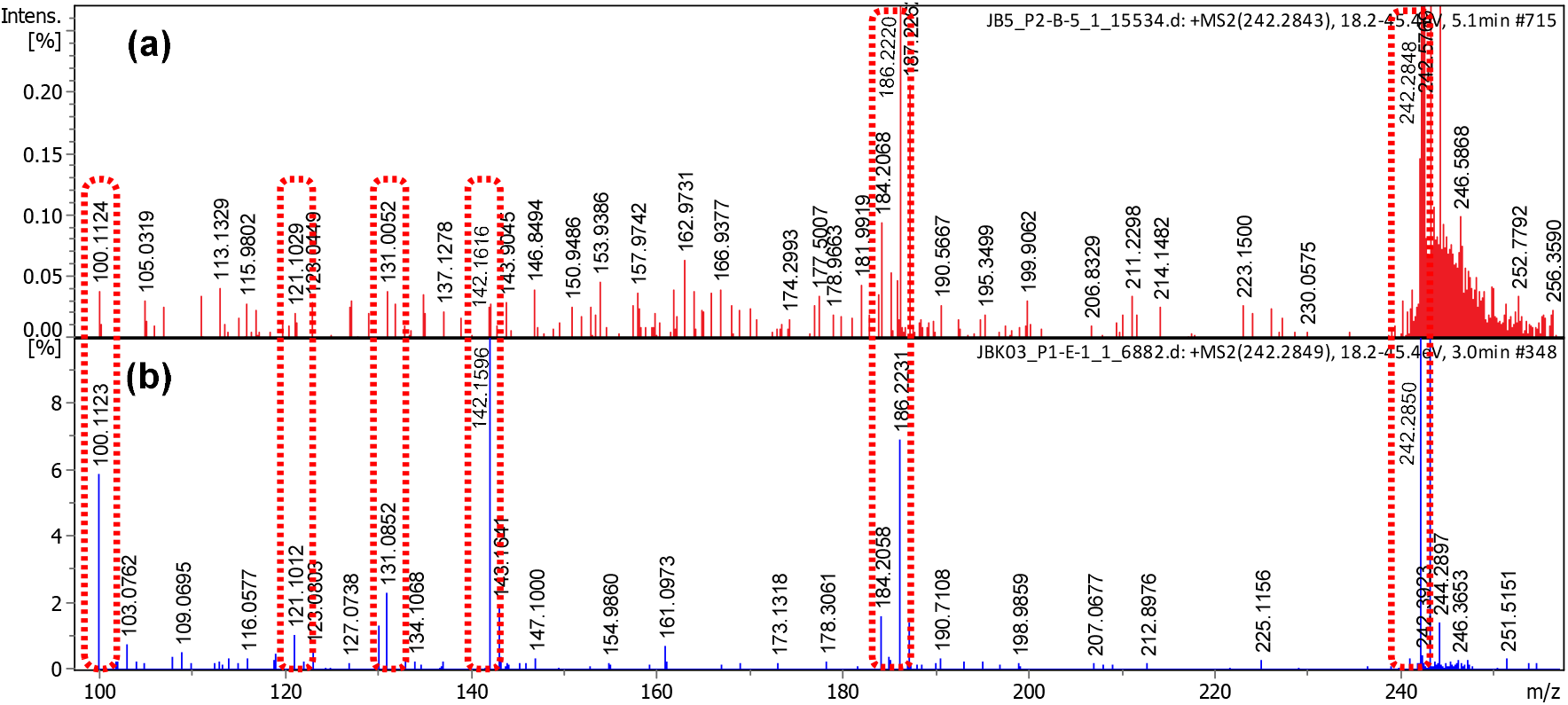
Tandem mass spectra of N-butyldodecan-1-amine (NBDA) in (a) pure compound standard and (b) rice genotype (Koshihikari) Similar fragmentation patterns between both mass spectra have been highlighted by boxes, confirming the presence of NBDA compound in the rice exudates.

